# Rapid Detection of Active Coronavirus Infection by Lateral Flow Test Strips: A New Approach to Distinguish Replicating Viruses from Non-Replicating Viruses

**DOI:** 10.1101/2024.07.18.604218

**Authors:** Darnell Davis, Caitlin H. Lamb, Cameron Myhrvold, Hemayet Ullah

**Affiliations:** Department of Biology, Howard University, Washington, DC; Department of Molecular Biology, Princeton University, Princeton, New Jersey, 08544, USA; Omenn-Darling Bioengineering Institute, Princeton University, Princeton, New Jersey, 08544, USA; Department of Chemical and Biological Engineering, Princeton University, Princeton, New Jersey, 08544, USA; Department of Chemistry, Princeton University, Princeton, New Jersey, 08544, USA

## Abstract

This manuscript describes the development of an alternative method to detect active coronavirus infection, which targets negative-sense RNA, a product of active viral replication. Few diagnostic methods are capable of discriminating between replicating and non-replicating viruses, complicating decisions related to quarantine and therapeutic interventions. We propose strand-specific nucleic acid diagnostics as a means of distinguishing between active and inactive RNA virus infections and prototype a CRISPR-based lateral flow assay that specifically detects replicating coronaviruses. Such a paradigm in diagnostics could guide more effective public health measures to curb the spread of SARS-CoV-2 and other single-stranded viruses.

## Introduction

The COVID-19 pandemic, caused by the rapid spread of SARS-CoV-2, was significantly exacerbated by delays in the development of rapid and affordable diagnostics [1–3]. Although traditional diagnostic methods like quantitative polymerase chain reaction following reverse-transcription (RT-qPCR), rapid antigen tests, and sequencing were critical to understanding the early spread of the outbreak, they notably lacked an ability to distinguish active SARS-CoV-2 infections from inactive remnants and were fundamentally unable to quantify virus replication dynamics [4]. Positive-strand RNA viruses, including coronaviruses, can persist even after active infection has resolved [5]. This persistence can limit the clinical utility of PCR-based tests that are strand-agnostic and will have similar readouts when detecting replicating virocells as for newly exocytozed virions and residual and/or neutralized viral particles that can remain after an infection has passed [6–8]. Antigen tests have lower sensitivity and specificity compared to nucleic acid-based tests due to our inability to amplify antigens and are similarly fundamentally limited at distinguishing between replication-competent particles and persistent fragments. Meanwhile, culture- and some sequencing-based methods can elucidate infectivity and viral replication dynamics but are made irrelevant in their deployability for informing immediate clinical decisions by cost and duration.

Severe through asymptomatic cases of COVID-19 have been widely reported to continue harboring or shedding detectable viral sequences and proteins long after infection, but the literature has yet to resolve whether these fragments are indicative of replication or are simply remnants of infection. Understanding viral shedding patterns and the duration of infectiousness is crucial for effective disease management and public health interventions. Owusu et al. found that while individuals may test positive for SARS-CoV-2 for extended periods, the presence of viral RNA does not necessarily indicate infectiousness [34]. The finding suggests that patients with mild to moderate COVID-19 are unlikely to be infectious beyond 10 days after symptom onset, despite potentially continuing to test positive [34]. This distinction between viral detection and infectious potential has important implications for isolation policies, patient care, and the management of disease spread in communities

A prominent publication reported persistent diffuse viral replication months after symptom onset in COVID-19 patients [9]. This was evidenced by the detection of sub-genomic viral RNAs in autopsied tissues and the replication of SARS-CoV-2 in Vero E6 cell cultures seeded with diverse autopsied tissues [9]. However, it remains unclear how these findings from fatal cases relate to less severe infections. On the other hand, many investigators attribute the long-term presence of the SARS-CoV-2 to non-replicating viral fragments [10–13]. Biopsies of patients with myocarditis showed the presence of SARS-CoV-2 proteins and RNAs up to 18 months after infection [10], although the viability of these particles was not assessed. One study found fragmented SARS-CoV-2 RNA and viral proteins in non-apoptosing non-classical monocytes of a patient 15 month post-infection without any indications of active replication [14]. Even in asymptomatic cases, viral sequences can be pervasive but may not necessarily be reason for concern in the clinic. SARS-CoV-2 viral RNA was detected in the tonsils of 20% of 48 children displaying no signs or symptoms of COVID-19 prior to a tonsillectomy [15]. In another study, some patients with negative PCR results who underwent gastric and gallbladder surgery more than a year after infection stained positive in SARS-CoV-2 nucleocapsid-specific immune-histologies [16], although it remains unclear whether such viral remnants would pose any risk to surgical staff because cultures were reportedly not feasible. Culture-dependent methods are the gold standard for determining viral replication competence, and by inference, infectivity, but they are low-throughput and require several days to perform by trained staff; a fact that would preclude their use in preventing transmission during solid organ transplant, or could lead to ill-informed guidelines for quarantining. A new, simpler method is urgently needed for determining if a person is harboring actively replicating virus to infer infectivity.

Here, based on our earlier work, we propose and show the effectiveness of a detection system that can directly target negative-sense viral RNA, produced only during replication. In contrast to positive-stranded fragments, these RNAs should not remain after infection has passed, thereby distinguishing replicating virus from non-replicating virus [17–19]. We show that the replicating mouse coronavirus, a model for SARS-CoV [20–22], continues to produce both its genome, positive-sense RNA (+ssRNA genome), and antigenome, negative-sense RNA (-ssRNA antigenome), throughout the infection process in cells [5, 17]. However, once replication has abated, we find that only the +ssRNA genome is still detectable from viruses in host cell-depleted culture media, indicating that -ssRNA antigenome may serve as a marker for viral replication competency and may thus indicate infectiousness.

Since we first proposed the idea, several groups have also shown the effectiveness of targeting the -ssRNA antigenome of the coronavirus as a means of detecting actively replicating viruses [19, 23–26]. However, a lack of user-friendly strand-specific tests prevent widespread adoption of this paradigm in diagnostics. We therefore prototype a user-friendly CRISPR-based lateral flow assay to overcome such shortcomings in the market of available detection technologies. Although our primary objective in this study does not directly address the persistence of replication-competent particles, the distinction between active infection and non-replicating infection afforded by stranded diagnostics can nevertheless inform containment strategies. Assuming active production of the antigenome increases the output of infectious particles, such a detection system could allow myriad advantages, ranging from more informed quarantine policies to effective therapeutic decisions.

## Results

### Use of qPCR to Detect Negative-sense RNA from Mouse Coronavirus MHV-A59

As a model system for coronavirus replication, we used murine hepatitis virus strain A59 (MHV-A59) harboring an eGFP fluorescent tag inserted by replacing its ORF4 pseudogene [27]. MHV has been widely used to model the Coronaviridae family of enveloped positive-stranded RNA viruses, and the MHV-A59 strain is the prototype murine betacoronavirus [28, 35]. We infected mouse 17CL-1 fibroblasts with the reporter virus and used qPCR to detect MHV sequences isolated from the infected cells two hours post-infection, according to the protocol depicted in Figure 1A. We used strand-specific cDNA primers to generate positive- and negative-strand cDNA libraries by reverse transcription (RT) and amplified an approximately 600 bp-long PCR product from the respective cDNA pools, alongside an actin control. Though the cDNAs were not generated with actin or oligo dT primer, actin-specific amplification was observed, possibly due to self-priming or DNA contamination in RNA samples. While not used for transcript quantification, these actin bands serve as a quality control check for strand-specific cDNAs. We found that the negative strand was more readily amplified from infected cell extracts than was the positive strand (Fig. 1B); this may be the result of strand-specific biases in RT efficiency or may imply more negative-strand synthesis immediately after infection. Nevertheless, we could reliably detect both the negative- and positive-strands in our model culture system.

**Figure 1:**
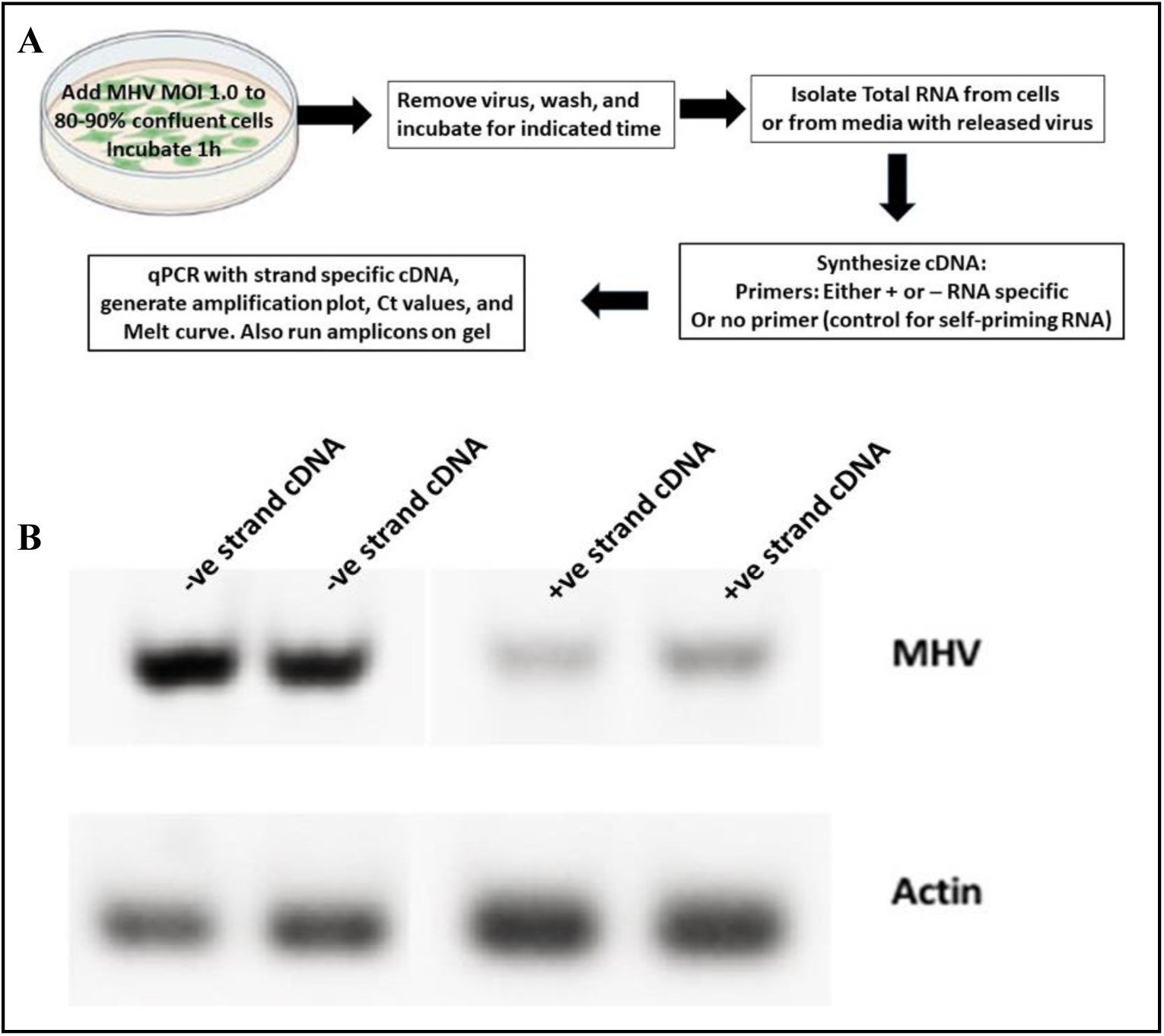
A. Flow chart depicting the experimental steps for detecting the MHV-A59 +ssRNA or -ssRNA strand. B. RT-PCR of the -ssRNA strand (lane 1 and 2 in duplicate) and the +ssRNA strand after 2h of infection in 17CL-1 cells with MOI 1.0 of MHV-A59.

Because antigenome production is necessary to sustain viral replication, we tested whether we could detect the negative strand continuously when active infection was underway. We isolated RNA samples from different time points post-infection, until almost all host cells were depleted by viral replication. Fig. 2 shows representative stages of the cells from which total RNA was isolated. By day 3 post-infection with an MOI of 1.0, almost all the host cells were depleted due to the virus’ cytopathic effect (Fig. 2C). In all cell-extracted RNA samples taken throughout the infection, we detected both positive and negative ssRNA strands (Fig. 1B). We also tested whether both strands could be detected in RNA samples isolated from cell-depleted media collected 72 h post-infection and included a primer-free cDNA synthesis control for the same RNA samples to rule out reverse transcriptase carry-over or RT “self-priming” [29]. By 24 h post-infection, the negative strand was substantially less detectable than the positive strand of the virus; a decrease that was further accentuated by 72 h post-infection, as the negative strand approached background detectability (Fig 2H). This establishes that the -ssRNA antigenome is most abundant when the virus is capable of replicating in host cells, and gradually reduces to undetectable levels as actively replicating virocells die off and infection-naïve cells are lost. Meanwhile, +ssRNA stranded viral RNAs increased in abundance up to the 72 h time point, representing a combination of virions and residual genomic fragments. Thus, positive-stranded viral RNA can be detected from both replicating and non-replicating environments, while replication is necessary for the negative strand to be detected. We normalized the samples using the respective cDNA concentrations taken by nanodrop (Supplementary S2A).

**Figure 2:**
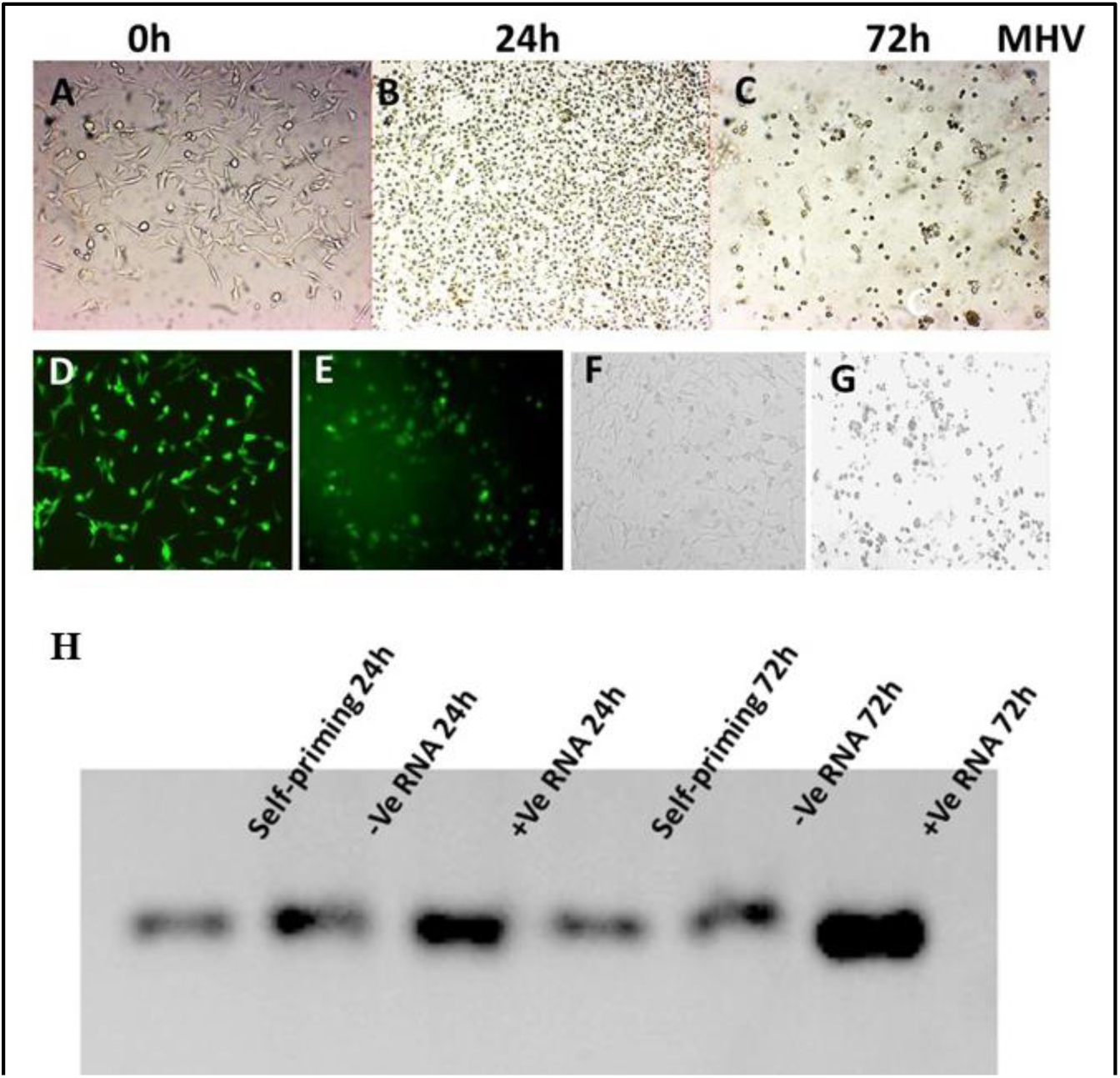
Developmental stages of the host cells(17CL-1) post infection with the MHV-eGFP-A59. The Cytopathic effect is completed by 72h post infection. Lower panel (D, E) depicts the visual effect of the GFP tagged virus infection showing intense fluorescence in the cells with incipient cytopathic effect and panel F and G show the same fluorescence cells in bright light. H. RT-PCR based PCR product targeting the +ssRNA or the -ssRNA strand. Lane 1 and 4: 24h post infection with self-priming cDNA; Lane 2 and 5-24h and 72h post infection with negative RNA specific cDNA samples, and lane 3 and 6-24h and 72h post infection with positive RNA strand specific cDNA samples. Samples were normalized to the concentrations of each of the cDNAs.

To better evaluate the relative abundances of these RNAs, the same cDNA from 72h post-infection were input into qPCR reactions (Table 1 & Fig. S2).

**Table 1:** Serial Dilutions of no primer (Self-amplification), -ve and +ve ssRNA strand specific cDNAs based Real-Time PCR results. The 2^-ΔΔCt^ method was used to calculate fold changes using the No Template Control as base line control. The mean Ct values were calculated from three technical replicate samples and the corresponding Standard Error of the mean (SEM) were calculated from the same samples.

|  | Mean<br>Ct (0.1<br>MOI) | SEM | Fold<br>Change | Tm | Mean<br>Ct<br>(0.01<br>MOI) | SEM | Fold<br>Change | Tm |
| --- | --- | --- | --- | --- | --- | --- | --- | --- |
| **Self** | 17.83 | 0.048 | 1 | 82.41 | 21.52 | 0.148 | 1 | 82.46 |
| **-Ve** | 17.75 | 0.099 | 1.05 | 82.42 | 21.34 | 0.046 | 1.12 | 82.31 |
| **+Ve** | 15.82 | 0.072 | 4.0 | 82.31 | 19.61 | 0.116 | 3.76 | 82.27 |

In concordance with the RT-PCR gels, the viral amplicon in the -ssRNA cDNA pool had an indistinguishable abundance from that of the primer-free cDNA control (experimental noise), while the same amplicon was much more abundant in the positive RNA-specific cDNA sample. Collectively, these data confirm that +ssRNA stand may remain abundant whether or not virus has stopped actively replicating, whereas negative-strand viral RNAs can only be detected above background levels when the virus is replicating.

### Using Cas13a to distinguish -ssRNA antigenome and +ssRNA genome of Mouse coronavirus

We next wanted to determine whether it would be feasible to develop a stranded nucleic acid detection assay to distinguish between samples containing only viral particles and those coming from cells with actively replicating viruses. We designed two *Leptotrichia buccalis* (*Lbu*)Cas13a guide RNAs that detect the 5’ region of the MHV-A59 genome’s reverse complement using ADAPT for Cas13a guide activity predictions in addition to NUPACK [33, 36] for RNA ensemble characteristics and minimum free energy predictions (see Materials and Methods). Our guides were in the same sense as the genome and had no significant pairwise local alignments with its complement, and so would not have any biological reason to detect the sense strand of the genome, while being designed to have perfect complementarity to its complement. We combined these guide RNAs with the enzyme and a short RNA reporter (an RNA-tethered fluorophore-quencher pair) in a fluorescent assay to detect the negative strand, wherein target recognition is detected as fluorescence of cleaved RNA reporters (Fig. 3A). As expected, *in vitro* transcribed RNAs representing the genome from an amplified region at its 3’ end was not detected, while samples containing its complement in the same amplified section could easily be distinguished from a negative control within minutes of starting the assay, with robust detection at 90 minutes (Fig. 3B and 3C). This assay clearly distinguishes the -ssRNA antigenome from the +ssRNA genome without requiring thermal cycling.

**Fig. 3:**
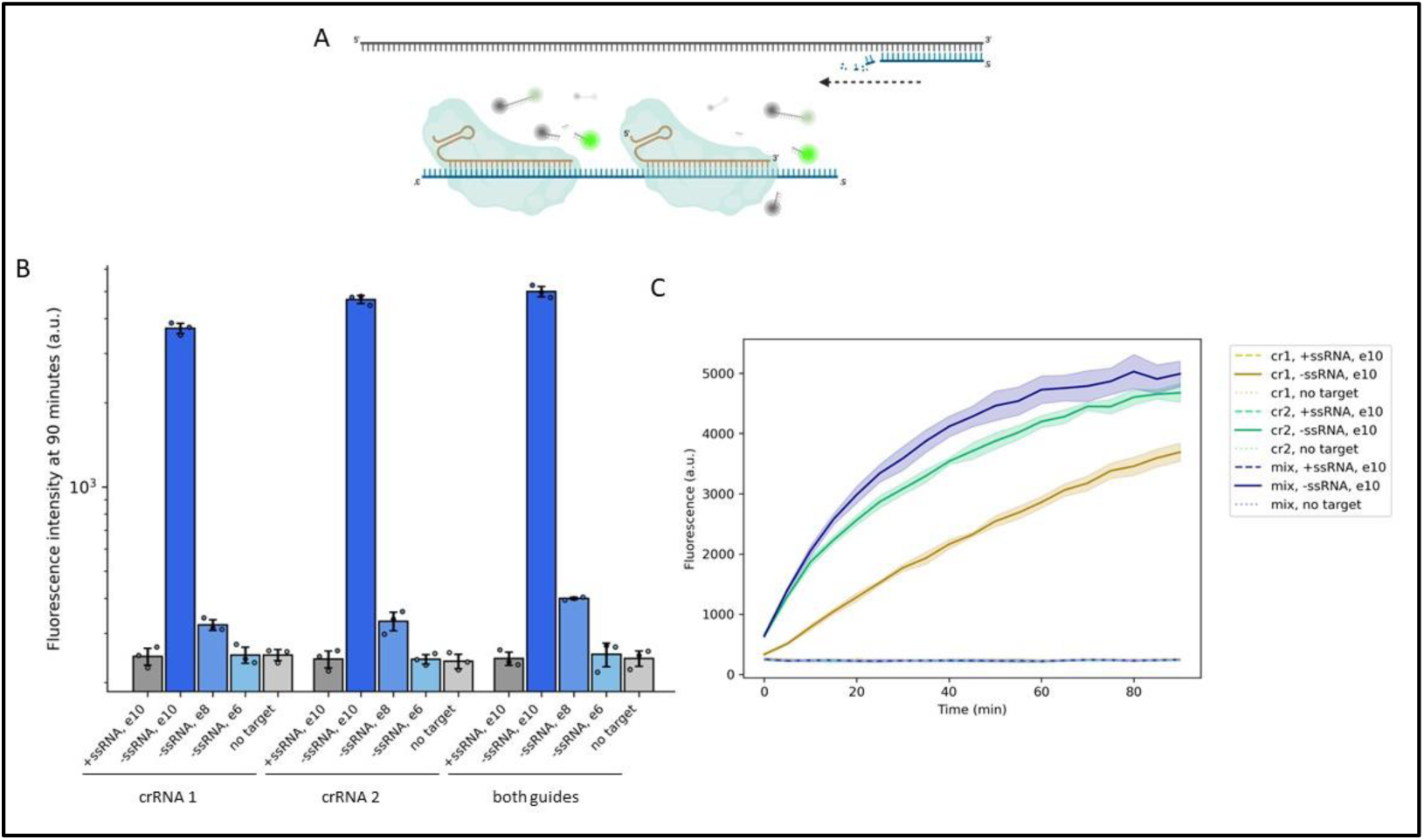
A. Non-promiscuous detection of stranded coronavirus RNA sequences *in vitro*. A) Schematic of stranded detection, wherein Cas13 (cyan) is guided by crRNAs (orange) sequence-specifically targeting the 5’ region of the replicating RNA genome (-ve sense, blue). Upon recognition of the replicating genome, the enzyme’s nuclease becomes active, cleaving nearby reporter RNAs and releasing fluorophores from quenchers. Doodle created using BioRender B. Quantitation of fluorescence after 30 minutes of incubating Cas13 with combinations of guide RNAs in the presence of purified RNA representing the +ve sense MHV genome (+ssRNA), different concentrations of RNAs representing the replicating genome template (-ssRNA), or no target. All concentrations are reported as RNA copies/µL in the sample before this was added to the detection master mix. e10 represents 10^10^ copies/µL, or a total of 1.5 X 10^10^ copies in the final 15µL reaction. Individual points represent three technical replicates per condition, while bars represent means ± standard deviation. C. Fluorescence curves over 90 minutes for the same conditions as in B. Lines and solid fills represent means ± standard deviation for three technical replicates per condition, taken at 5-minute intervals.

### Development of an LFT Targeting the Negative Strand

With the ultimate goal of designing a more user-friendly assay from the same CRISPR platform, we next moved to Lateral Flow Tests (LFTs) and opted to include additional guides to validate the assay’s sequence agnosticism. We designed two additional guide RNAs to detect the 5’ region of the MHV-A59 -ssRNA antigenome and two additional guide RNAs targeting the +ssRNA genome as a control. After incubating *Lbu*Cas13a with the respective guide RNAs, the ribonucleoprotein complex was added to RNA isolated from 17CL-1 cells 4h or 72h post-infection alongside a biotinylated RNA reporter. When using RNA (100 ng) from non-replicating viruses released in the media, the test band was only visible when using +ssRNA genome-targeting guide RNAs (Fig. 4A, left panel), supporting the strand-specificity of this assay. Conversely, when using RNA from replicating viruses within the host cells, both the genome- and antigenome-specific assays showed positivity, indicating the presence of both forward and template genomes at this stage of replication (Fig. 4A, right panel). Note that the RNA from replicating viruses was isolated 4 hours post-infection, and more than 99% originated from host cells. The identification of both the viral +ssRNA and -ssRNA strands while containing low amounts of viral RNA compared to the host RNA in the original sample indicate that this LFT system is reasonably sensitive in addition to being specific (left panel). Additionally, we confirmed the LFT results using a fluorescent RNA probe (see S1 Table for sequence) as above on 72h RNA extracts from non-replicating viruses (Fig. 4B), confirming a strand and concentration-dependent detection of genomic RNAs by only the genome-targeting guide. The data of the results presented in the Figure 4B are also used to statistically validate whether the data generated from the use of +ve guideRNA, but not the -ve guideRNA, in the non-replicating RNA sample produce significant difference when compared to no RNA used samples. We found that both the 10 and 50 ng non-replicating RNA sample produce statistically significant data when only +ve guideRNA was used while use of the -ve guideRNA didn’t produce any statistically significant data when compared with no RNA samples with -ve guideRNA (Fig. S3). The heatmap of the signals (Fig. 5) shows that over the 100 minutes of incubation period, it is only the +ve guideRNA sample (data not shown) with 10 and 50 ng non-replicating RNA produced steady increase of fluorescence indicating the binding of the +ve guide RNA to the targeted +ssRNA genome. As no increase in fluorescence was observed from the -ve guideRNA samples, compared to the no RNA samples, it can be concluded that the non-replicating RNA samples didn’t have the targeted -ssRNA genome in the sample RNA.

**Fig. 4:**
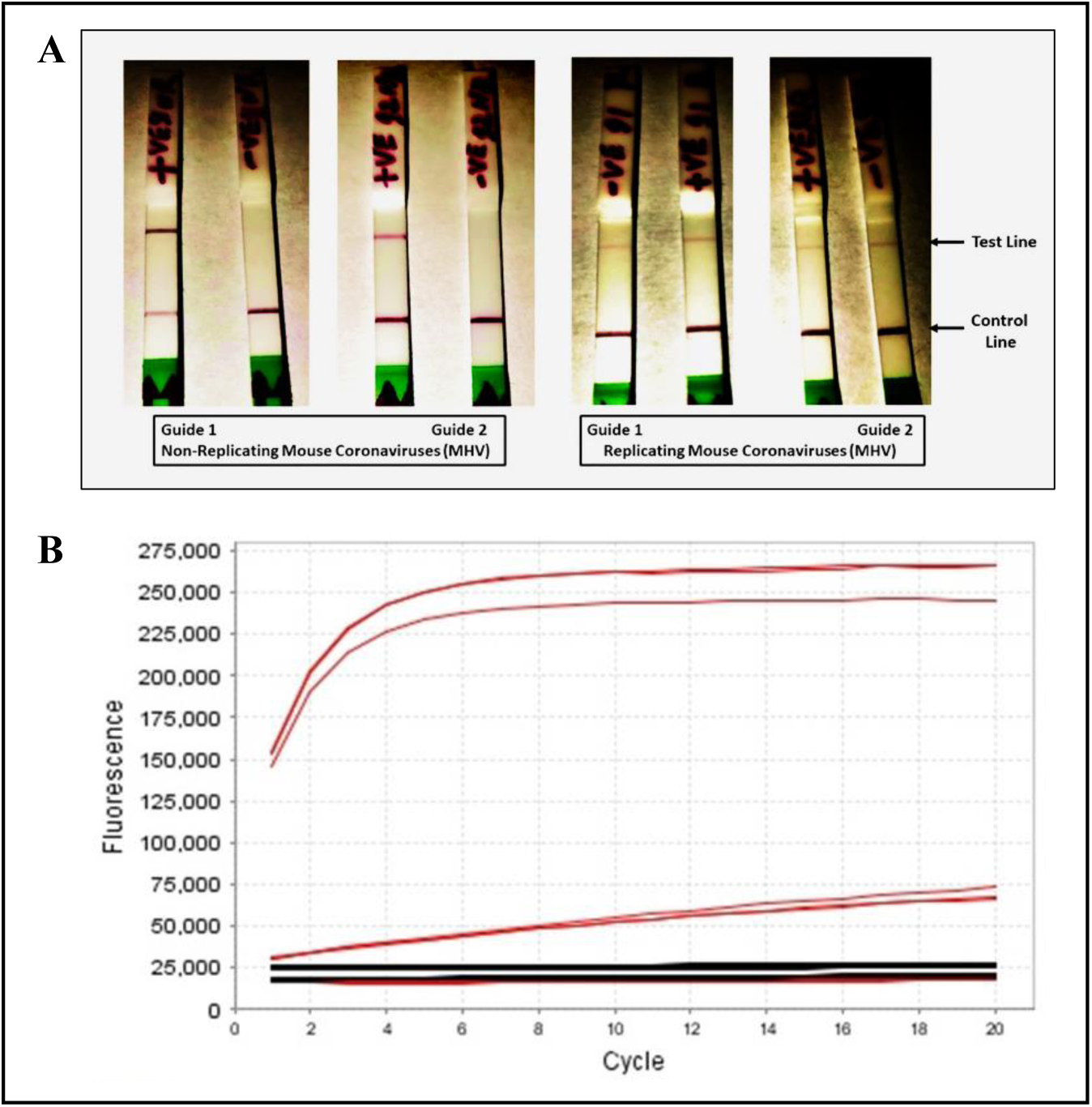
Representative lateral flow test results depicting detection of non-replicating and replicating coronavirus RNA samples. A. Right Panel: Results from replicating virus RNA samples demonstrate positive test bands in both the positive and negative ssRNA strand-specific test strips (the contrast was enhanced to clearly show the bands-reason in the text). Left Panel: Replicating virus RNA showing only positive strand specific band while negative strand specific band were absent. B. Fluorescent probe-based detection of the positive strand from the non-replicating virus. Positive strand specific guide RNA (red line) cleaved FAM signal is depicted by increasing fluorescence identity in triplicate samples. The negative strand specific guide RNA (black line) based signal does not show any increasing fluorescence signal (in triplicates). No RNA sample is used as no-template control that shows a flat signal at around 23,000 fluorescent intensity signals similar to the negative strand specific signal intensity.

**Fig. 5:**
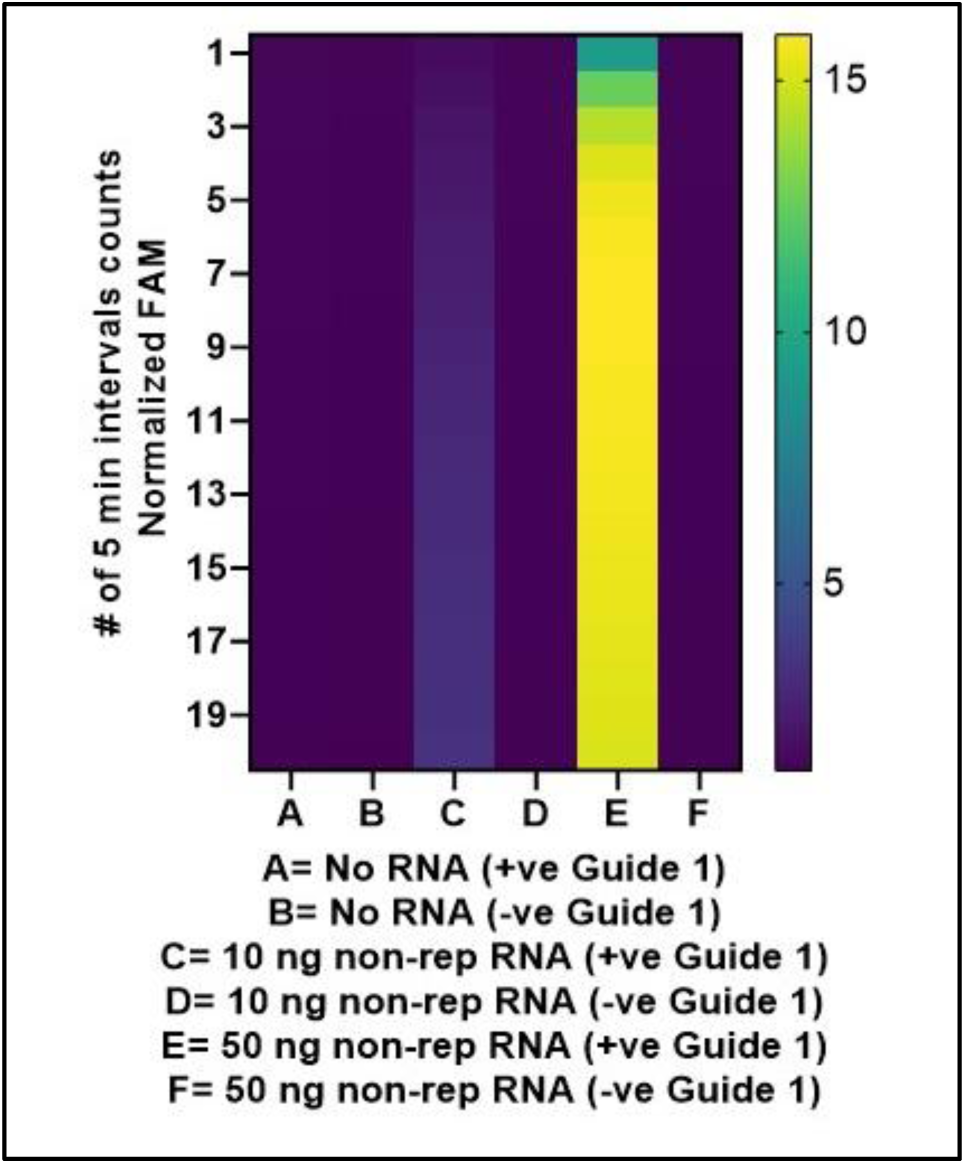
CRISPR-based fluorescent signal-based detection of positive viral RNA strands in non-replicating coronavirus samples. Heatmap of the FAM fluorescence representing every 5 minutes of 20 total readings for the samples stated in the figures.

Like the FAM signal data from the non-replicating RNA samples, we also plotted the median FAM fluorescence signal from the replicating virus RNA data for the 24 readings where each data point was computed as an average of three technical replicate samples. In spite of low viral RNA concentrations in the sample, both +ve (Fig. 6A) and -ve guide RNA (Fig. 6C) based FAM median data clearly showed statistically significant fluorescence when compared with corresponding no RNA samples. Similarly, the heatmap also showed steady increase in the FAM fluorescence over time for both the +ve guideRNA (Fig. 6B) and the -ve guide RNA (Fig. 6D) based fluorescent signals. The test strips also showed similar test band indicating the presence of both the -ve and +ve RNA genome within the sample (Fig. 4A). Note that the heatmap clearly can reveal that the sample contained very low level of viral RNA, hence, the increase in the fluorescence was not as robust as in the non-replicating RNA samples with the +ve guideRNA.

**Figure 6:**
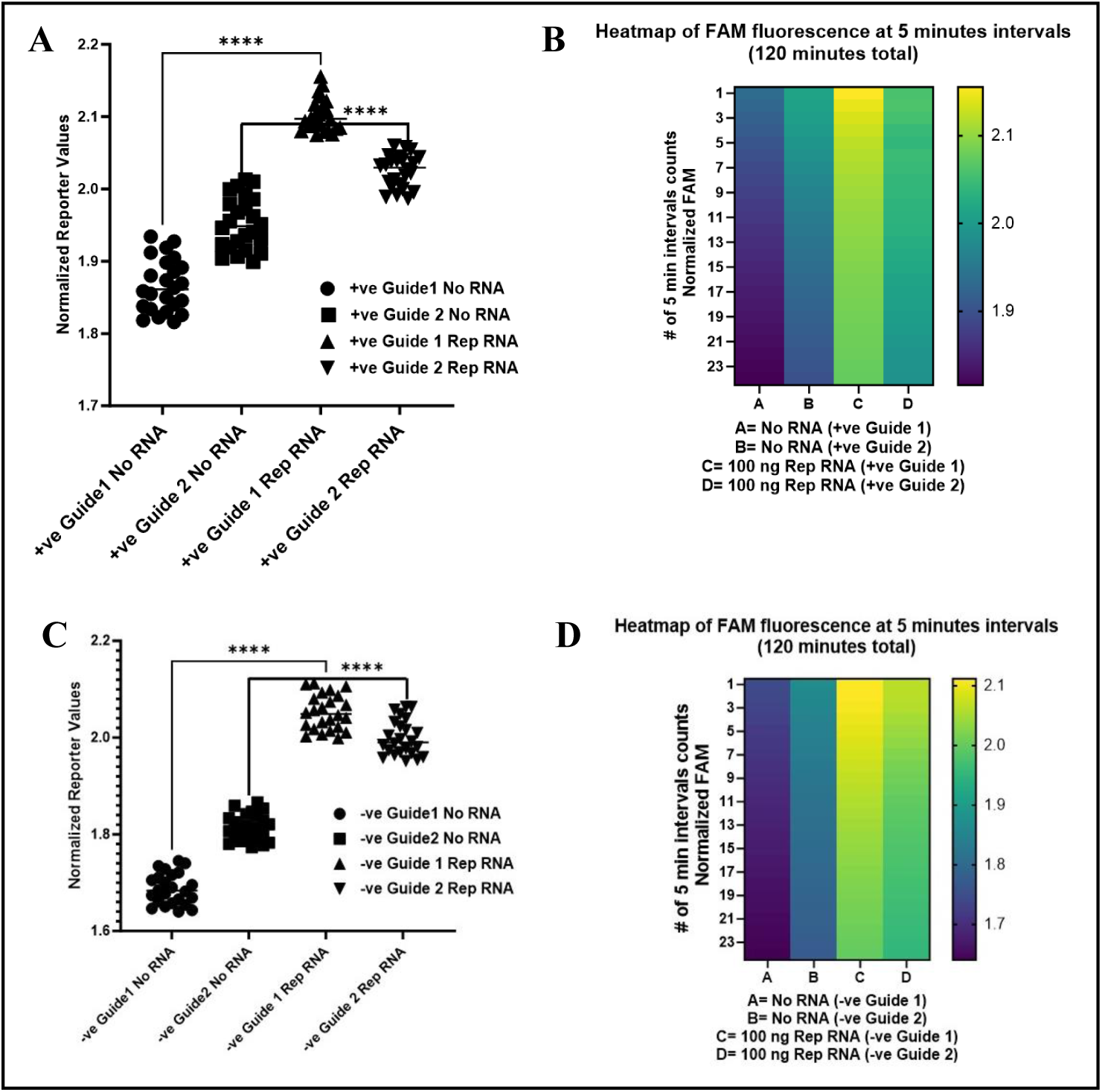
CRISPR-based fluorescent signal-based detection of positive viral RNA strands in replicating coronavirus samples. A. The median signal from the cleaved FAM fluorescent signals every 5 minutes in total 120 minutes are plotted. Each data point represents mean of triplicate samples. The significance of the pairwise comparison of median signals is indicated as significant (**** p<0.0001) or as not significant (ns). B. Heatmap of the FAM fluorescence representing every 5 minutes of 24 total readings for the same samples represented in panel A. C. The median signal from the cleaved FAM fluorescent signals every 5 minutes in total 120 minutes are plotted. Each data point represents mean of triplicate samples. The significance of the pairwise comparison of median signals is indicated as significant (**** p<0.0001) or as not significant (ns). D. Heatmap of the FAM fluorescence representing every 5 minutes of 24 total readings for the same samples represented in panel A.

The plate reader was also used to measure the end-point fluorescent signal of the samples (diluted 1:5 with water) with the replicating viral RNA targeted by the +ve or by the -ve guideRNA. The results as depicted in the supplementary data (S2), showed statistically significant differences in both the +ve and the -ve guide RNA based FAM fluorescence compared to the corresponding no RNA samples. It is evident that the extremely low level of viral RNA in replicating virus samples isolated from host cells prevented strong FAM signals compared to the signals from non-replicating sample. This result, combined with the strong separation of signals between RNA from replicating and non-replicating viruses, suggests potential utility in distinguishing between these viral states. These findings highlight the utility of the lateral flow test in discriminating between non-replicating and replicating coronavirus infections based on strand-specific detection.

## Discussion

The COVID-19 pandemic has had a profound impact on the world, with the virus spreading rapidly and causing significant illness and death since it started in 2020. To counter the pandemic, researchers and healthcare professionals have been developing new tools and technologies for the detection and diagnosis of the virus. However, the inability to distinguish replicating viruses from non-replicating viruses limits the utility of the current detection methods. We described the development of an alternative active coronavirus detection system that uses the -ssRNA antigenome as a marker for the presence of a replicating virus.

One of the key advantages of using the -ssRNA antigenome as a target for detection is its ability to distinguish between active and non-active infections. By targeting the -ssRNA antigenome, this detection approach should be able to detect only those individuals who are currently infected with actively replicating virus, and thus are capable of transmitting it to others. This is an important distinction, as individuals who have recovered from the virus may still test positive using other detection methods targeting the +ssRNA genome, but that may not necessarily indicate infectiousness, as remnant/residual virus genome fragments can yield positive results in the absence of any replication.

Furthermore, the presence of a negative strand is a clear indicator of active infection. If an individual’s sample does not contain the negative strand, this indicates that they are not currently infected with any actively replicating virus and do not need to be quarantined. [11, 32]. This can help reduce the burden on healthcare systems and prevent the unnecessary isolation of individuals.

Another key strength of this approach is its high sensitivity and specificity. By targeting the -ssRNA antigenome, our system was able to accurately detect the presence of the virus, even at low levels. This is particularly important in the early stages of the disease, when the amount of virus in an individual sample may be low. Additionally, the use of the -ssRNA antigenome as a target allows for the specific detection of coronavirus, reducing the potential for false positives.

Additionally, the use of the -ssRNA antigenome of coronavirus as a marker for replicating viruses may be useful for monitoring the effectiveness of antiviral therapies. By measuring changes in the levels -ssRNA antigenome/strand over time, clinicians can determine whether a particular treatment effectively reduces the number of replicating viruses in the body. This information can be used to guide treatment decisions and optimize patient outcomes.

In our continued efforts to accurately detect and differentiate active viral infections, we have also developed a prototype lateral flow test capable of selectively identifying replicating viruses. Building upon the RT-PCR results above to distinguish replicating from non-replicating viral RNA, this novel lateral flow test represents a significant advancement for rapid, accessible testing. Lateral flow tests have emerged as powerful diagnostic tools during the COVID-19 pandemic, enabling widespread self-testing outside clinical settings. However, current LFTs cannot differentiate between an active, ongoing infection and merely the presence of residual, non-infectious viral particles. Our LFT overcomes this limitation by specifically targeting a unique molecular signature present only in actively replicating viruses.

The key advantages of our replication-specific LFT lie in its ability to provide a clear, unambiguous indication of active viral infection while avoiding false positives from past, resolved infections. By focusing the detection on the transient replication intermediates, our LFT effectively captures the dynamic state of viral propagation within the host. This not only improves diagnostic accuracy but also offers valuable insights into the infection stage and transmission risk. Unlike conventional LFTs that merely detect viral antigens or nucleic acids, our test distinguishes replicating from dormant viruses, a crucial distinction for informed decision-making and treatment strategies. With its user-friendly format and rapid turnaround time, this LFT can be readily deployed for large-scale screening, surveillance efforts, and personal testing, empowering individuals and communities with tailored, actionable information about their infection status. This innovative approach represents a significant advancement in diagnostic capabilities, particularly in the context of the current pandemic landscape. Unlike traditional diagnostic tests that may detect both active and inactive viruses, our LFT provides enhanced specificity, thereby minimizing the risk of false-positive results and enabling more accurate diagnosis of active infections.

The implementation of our LFT holds several key advantages over existing diagnostic modalities, particularly in the context of widespread testing initiatives and decentralized healthcare settings. By selectively detecting actively replicating viruses, our method offers rapid and reliable identification of individuals who pose an immediate risk of viral transmission. This targeted approach not only facilitates prompt implementation of infection control measures but also supports more effective allocation of resources for patient care and public health interventions. Furthermore, the simplicity and accessibility of lateral flow tests make them well-suited for point-of-care and at-home testing applications, offering greater convenience and flexibility for individuals seeking timely diagnosis and monitoring of their viral status. In essence, our LFT represents a pivotal advancement in diagnostic technology, poised to significantly enhance our ability to combat infectious diseases and mitigate the impact of pandemics on global health.

Development of this detection approach for active infection can have wide applications as a very large group of viruses belonging to many families are found to replicate through positive to negative RNA strand method. Along with coronaviruses, other positive RNA viruses that cause epidemic diseases in humans include encephalitis, hepatitis, polyarthritis, yellow fever, dengue fever, poliomyelitis, and common cold-causing viruses. Widespread use of the negative strand-specific detection system will not only address the most pressing issues of the current pandemic but will also present opportunities to address other disease-causing RNA viruses as well.

Though universal vaccination against the virus has occurred in some parts of the world, apprehensions of so called immune-escape-variants raise concern for a bleak future for the pandemic. The potential consequences of emerging variants are increased transmissibility, increased pathogenicity and the ability to escape natural or vaccine-induced immunity [2]. Several drugs with efficacy against the virus have been developed that are effective in reducing mortality and hospitalization. However, limited availability of drugs, contraindications, toxicity, rebound, etc. limit the widespread use of antiviral drugs [3, 4]. In addition, drug-escape-variants, like immune-escape-variants, can render existing treatments ineffective. All these issues make the effective management for the containment of the pandemic daunting, and the impact of the continued pandemic is causing widespread disruption to daily life around the world.

The use of the -ssRNA strand as a marker allows for the direct detection of replicating viruses, rather than relying on indirect indicators, such as the presence of viral proteins or antigens. This could result in a more sensitive and specific detection method. The use of the approach allows for real-time monitoring of virus replication, as the negative strand is produced during active viral replication. This could provide valuable information for disease management and control.

Overall, our findings support the establishment of the -ssRNA strand as a valuable target for active coronavirus detection. Further research is needed to evaluate the performance of this system about its sensitivity and specificity against different variants of SARS-CoV-2 and then testing on human samples, and to assess its potential for use in real-world settings. However, our results suggest that this approach has the potential to significantly improve prospects for the rapid identification of infected individuals and ultimately, in combatting the spread of the virus.

## Acknowledgements

The following reagents were obtained from BEI Resources (NIAID/NIH): Recombinant Murine Coronavirus MHV-A59 with Enhanced Green Fluorescent Protein (eGFP) (Cat# NR-53716), Murine 17Cl-1 Cell Line (derived from 3T3 cells) (Cat# NR-53719). We thank Owen Dunkley (Princeton University) for providing data and interpretations related to figure 3 as a proof of principle, in addition to substantive advice for improving certain sections of the manuscript.

## Materials and Methods

### Cells and Viruses

Murine 17CL-1 cell line (derived from 3T3 cells) was obtained through BEI Resources, NIAID, NIH, catalog number: NR-53719. The cells were maintained as monolayer cultures in Minimum Essential Medium (MEM; Sigma Aldrich-M 4655) containing 10% fetal bovine serum (FBS; Life Technologies), 100 IU/ml of penicillin, and 100 μg/ml of streptomycin (Both from Life Technologies) in a 37°C humified incubator supplemented with 5% CO2. MHV strain A59-eGFP, which expresses the Enhanced Green Fluorescent Protein (eGFP) inserted in place of the ORF4 gene, was obtained through BEI Resources (NR-53716).

### Infection

The 70-80% confluent 17CL-1 cells were infected with the MHV-A59-eGFP at 0.5 to 1.0 MOI. After one hour of adsorption, the media was removed, and the cells were washed three times with Phosphate-buffered saline (PBS). New MEM was added to the virus-treated cells and was incubated for the indicated duration.

### RNA isolation and RT-(q)PCR experiments

Viral RNA was isolated from media using the PureLink Viral RNA/DNA kit (Cat#12280) according to the manufacturer’s instructions (Thermo-Fisher, Carlsbad, CA). As detailed in the flow chart (Fig 1A), 500 ng of the total RNA from the media were used to synthesize strand-specific cDNA using the High-Capacity cDNA Reverse Transcription kit (Cat# 4368814) (Thermo-Fisher, Carlsbad, CA) and one of NegcDNA.F or PoscDNA.R (Table S1). The negative cDNA primer NegcDNA.F spans 30691 to 30713 bp of the murine hepatitis virus genome (accession #AY910861), while the positive RNA strand primer (PoscDNA.R) spans the region 31251-31273 bp of the murine hepatitis virus genome. Using both primers, a 582 bp long RT-PCR product was then amplified from the respective cDNA pools. As a loading control, mouse actin primers mActin.F and mActin.R were used to amplify a 154 bp long PCR product. RNA Isolation from Infected Cells:

Total RNA was isolated from samples using TRIzol reagent (Invitrogen, Carlsbad, CA) following the manufacturer’s protocol with minor modifications. Briefly, samples were homogenized in 1 mL of TRIzol reagent per 50-100 mg of infected cells. The homogenate was incubated at room temperature for 5 minutes to permit complete dissociation of nucleoprotein complexes. Chloroform (0.2 mL per 1 mL of TRIzol) was added, and the mixture was shaken vigorously for 15 seconds. After 3 minutes of incubation at room temperature, the samples were centrifuged at 12,000 x g for 15 minutes at 4°C. The aqueous phase containing RNA was carefully transferred to a new tube. RNA was precipitated by adding 0.5 mL of isopropanol per 1 mL of TRIzol used. Samples were incubated at room temperature for 10 minutes and then centrifuged at 12,000 x g for 10 minutes at 4°C. The RNA pellet was washed once with 75% ethanol, air-dried for 5-10 minutes, and resuspended in RNase-free water. RNA concentration and purity were determined using a NanoDrop spectrophotometer (Thermo Scientific, Waltham, MA).

qPCR: Quantitative PCR was performed on nanodrop-normalized sample inputs using the SYBR Select master mix (Cat# 4472908, ThermoFisher, Carlsbad, CA) on a StepOnePlus Real-Time PCR system (Applied Biosystems) starting with a single two-minute 50°C cycle (UDG inactivation) followed by two minutes at 95°C, 40 cycles of 95°C for 15 s and 60°C for 60 s, in addition to a melting curve cycle.

### Synthesis of strand-pure RNA targets for the CRISPR assay

cDNA of the 3’ end of the MHV-A59 genome from above, was amplified by PCR using MHV_rvRNA_FW and MHV_rvRNA_RV to template –ssRNA synthesis or MHV_fwRNA_FW and MHV_fwRNA_RV to template +ssRNA synthesis (Table S1). cDNA was amplified by 25 µL PCR using Q5 polymerase (NEB #M0492) under manufacturer-suggested conditions with 25 cycles of 98°C for 10 s, 66°C for 10 s, and 72°C for 30 s, and isolated using column purification (Zymo #D4013). RNA was expressed overnight in 30 µL T7 transcription reactions (NEB # E2050S, includes DNAse) with 3 ng of either DNA template, followed by DNAseI digestion for 30 minutes and bead-based RNA purification (Ampure A63987).

### CRISPR-based detection of stranded RNA sequences

Strand-specific CRISPR-Cas13a detection assays were performed using *Lbu*Cas13a and cognate guide RNAs designed to target non-repetitive elements at the 5’ end of the MHV-A59 genome’s reverse complement. Guide RNAs were selected by first generating a list of all possible candidates tiling the non-repetitive 537 bp region at the 5’ end of the replicating negative strand of the MHV-A59 genome between the amplification primers (NCBI Reference Sequence: NC_001846.1). These candidates were run through predictor_call.py using ADAPT (github.com/broad institute/adapt) [36] or their predicted activity in activating cleavage by *Lwa*Cas13a, an ortholog of the Cas13a protein being used in this study. We chose two candidates with high predicted Cas13 activity and minimal predicted undesirable RNA secondary structure in the spacer region using NUPACK with default parameters (NUPACK) [33], both targeting the negative-stranded replicating genome (Table S1).

15uL detection reactions were performed in 20 mM HEPES (pH = 8.0), 60 mM KCl, 3.5% PEG-8000, 1 U/µL Murine RNAse Inhibitor (NEB #M0314), 62.5 nM 6U_FAM_reporter (Table S1), 60 nM total crRNAs, 60 nM *Lbu*Cas13a (GenScript) and 14 mM Mg-acetate. 10% of the final reaction volume was sample; reported concentrations refer to the concentration of the target RNA diluted in RNAse-free water before the sample was input into the reaction. Reactions were mixed, covered, and incubated without shaking at 37°C in a Cytation5 plate reader, fluorescence curves were recorded using 20 nm-wide excitation and emission cutoffs, centered around 485 nm and 528 nm respectively. The templates, primers, reporter, and crRNA sequences are shown in table S1.

### CRISPR-based LFT development

Additional guide RNAs targeting the MHV-A59 positive or negative genomes were developed using the NCBI Reference sequence AY700211.1 (MHV-A59). The reference sequence in both the positive and negative sense were run through the ADAPT website (https://adapt.run/run/) [36]. As we were interested in direct detection of the positive or negative sense genomes, no penalty was added for primers, and the highest scoring spacers were used. The sequences of the 4 guide RNAs (two for the negative strand and two for the positive strand) and reporters are shown in table S1. The LbuCas13a Nuclease was purchased from the Genscript USA, Piscataway, NJ. The manufacturer’s protocol was used to assemble the crRNA and Cas13a complex. Briefly, 1 µl 10X Cas13a Reaction Buffer, 15 ng of strand-specific crRNA, 200 ng of GenCRISPR™ *Lbu*Cas13a Nuclease (Z03742) from 100 ng/µl stock, and DEPC-Treated Water (nuclease-free) was used to make a 10 µl reaction mix that was incubated for 10 min at 37^0^C.

For later fluorescent assays, indicated amounts of RNA from replicating or non-replicating viral samples were mixed with 10 µl of the ribonucleoprotein complex alongside 10 pmol of the 6UFAM reporter, DEPC-Treated water, and buffer for a 50 µl reaction, incubated at 37^0^C in a qPCR machine reading FAM fluorescence every 5 minutes, or equivalently in a Cytation5 plate reader as above.

For the LFTs, the same protocol was used with two modifications: instead of 6UFAM reporter, the FAM/Biotin Reporter was used. The assay mix (sample, crRNA/Cas13a, and RNA reporter) were incubated for 30 minutes before adding Hybridetect test strips (Milenia Biotec). The strips were allowed to develop for 5 minutes before photos were taken.

## Supplementary Figures

**Supplementary fig. 1:**
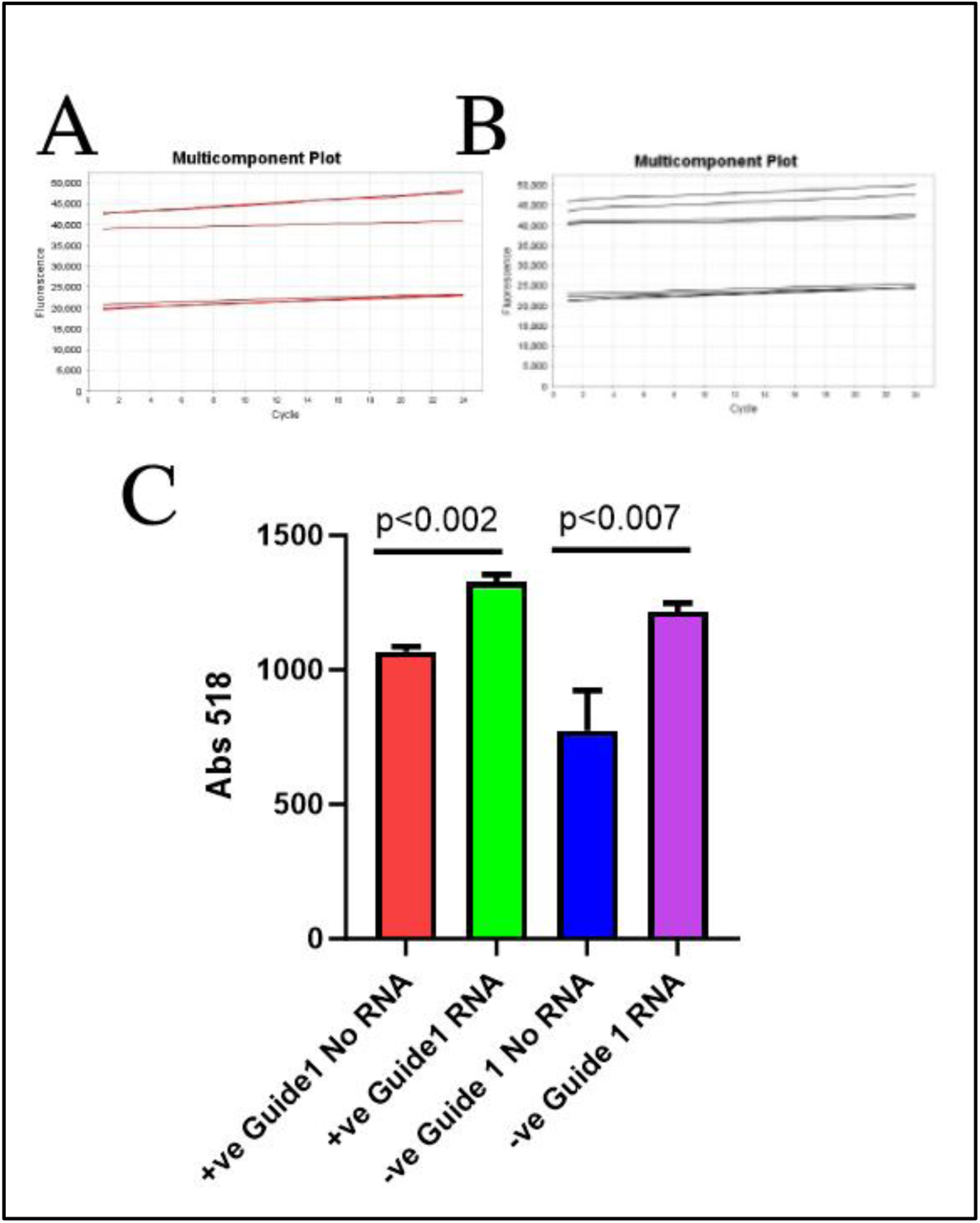
CRISPR-based fluorescent signal-based detection of negative and positive viral RNA strands in replicating coronavirus samples. A. Fluorescent signal from the cleavage of an RNA reporter by the LbuCas13a/guide RNA complex specifically bound to the positive (+) strand of the viral genome in a replicating virus sample. The accumulation of fluorescent signal over time indicates the presence of the positive sense genomic RNA strand. B. Fluorescent signal resulting from the cleavage of the RNA reporter by LbuCas13a/guide RNA complexes bound to the negative (-) strand of the viral genome in the same replicating virus sample. The increase in fluorescence confirms the presence of the negative strand replication intermediate. C. Quantification of the fluorescent signals from (A) and (B) measured in a plate reader with filters set at 494 nm excitation and 518 nm emission to detect the FAM reporter fluorescence over time. Sample with guide RNAs targeting the negative (-) strand is shown in purple, while that with positive (+) strand guides is in green. The increasing fluorescence in both sets of samples compared to the no RNA samples controls (Red for positive strand and blue for negative strand) provides orthogonal confirmation that the replicating virus contains both positive genomic RNA and negative strand replication intermediates. The graph represents signals from three technical replicates. Pairwise t-tests comparing each set of triplicate samples show statistically significant differences as indicated by the p-values provided above the respective bar graph pairs.

**Supplementary fig. 2:**
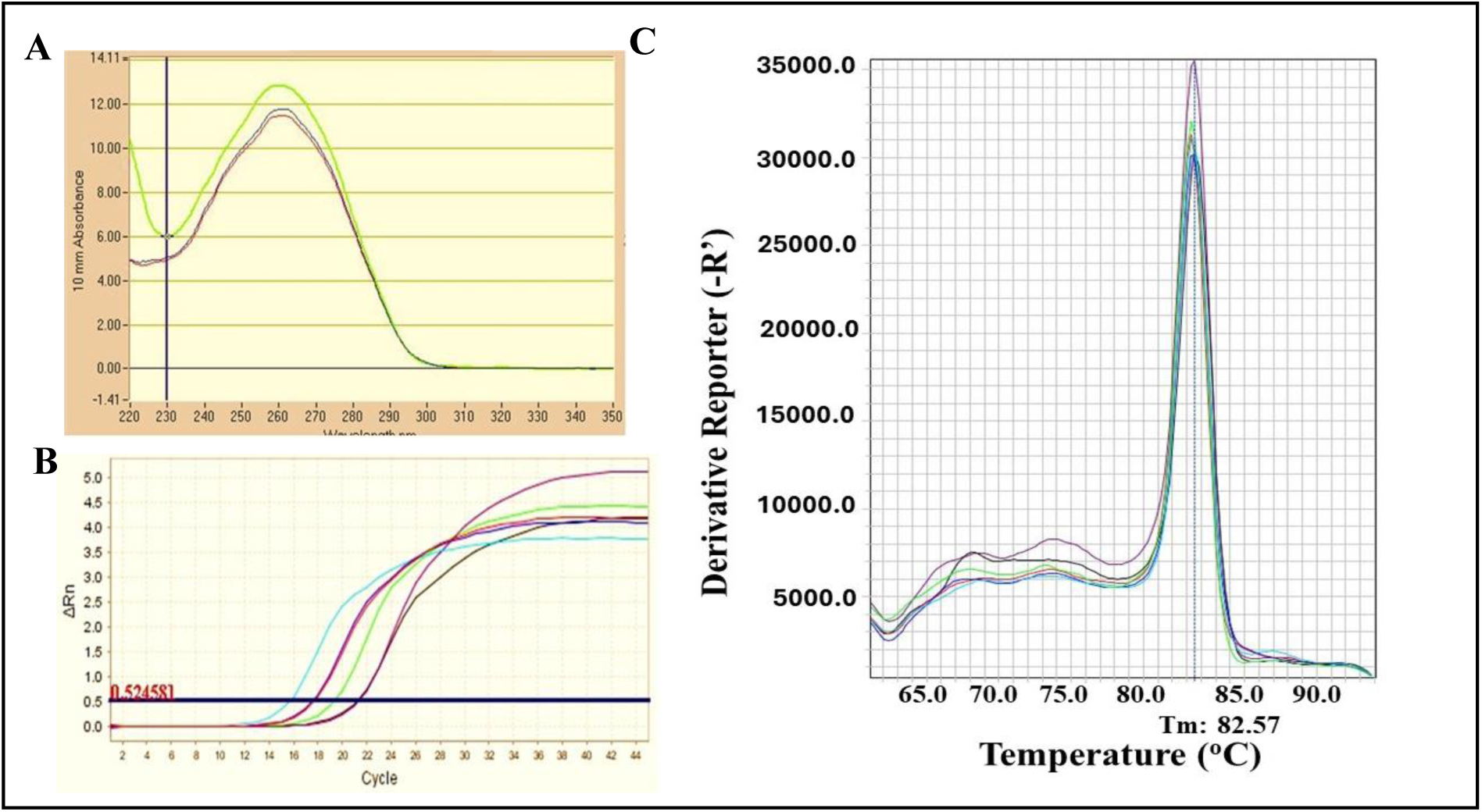
qPCR amplification plot of the negative and positive RNA strand specific cDNA samples. cDNAs from the 72h post infection samples were shown which was amplified at 0.01 and 0.1 dilutions of the cDNAs. Purple and black amplification plots in panel A represent 0.01 dilution samples from self-priming and negative-RNA strand specific cDNAs while green amplification plot represents the positive RNA strand specific cDNA sample. Red and blue amplification plots represent the self-priming and negative strand specific cDNA samples while the turquoise amplification plot represent the positive RNA strand specific cDNA sample at 0.1 dilution. Panel B shows the absorption spectra of all three cDNA samples. Panel C shows the melting curve of the amplified products in the panel A.

**Figure S3:**
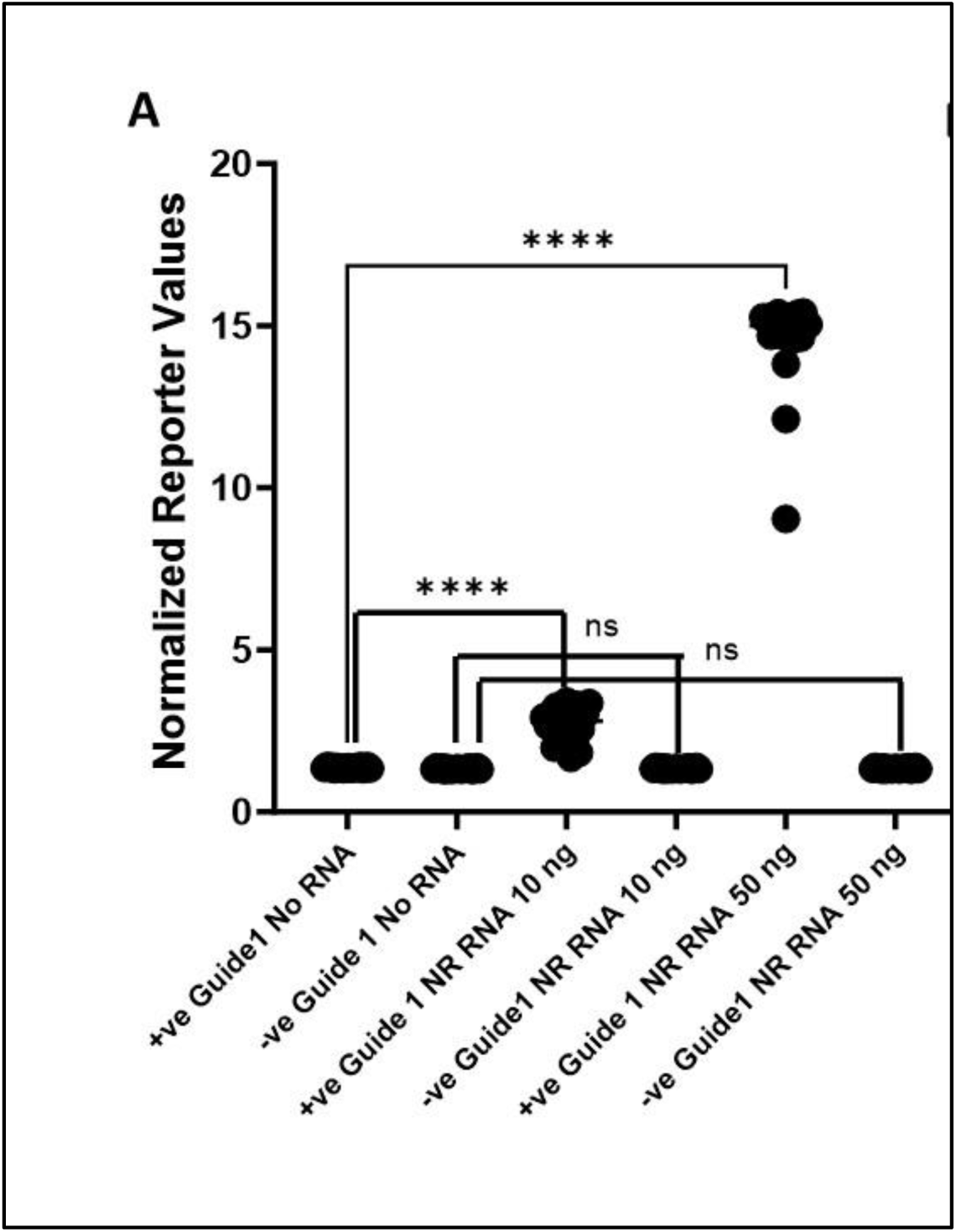
CRISPR-based fluorescent signal-based detection of positive viral RNA strands in non-replicating coronavirus samples. The median signal from the cleaved FAM fluorescent signals every 5 minutes in total 100 minutes are plotted. Each data point represents mean of triplicate samples. The significance of the pairwise comparison of median signals is indicated as significant (**** p<0.0001) or as not significant (ns).

**TABLE S1:**
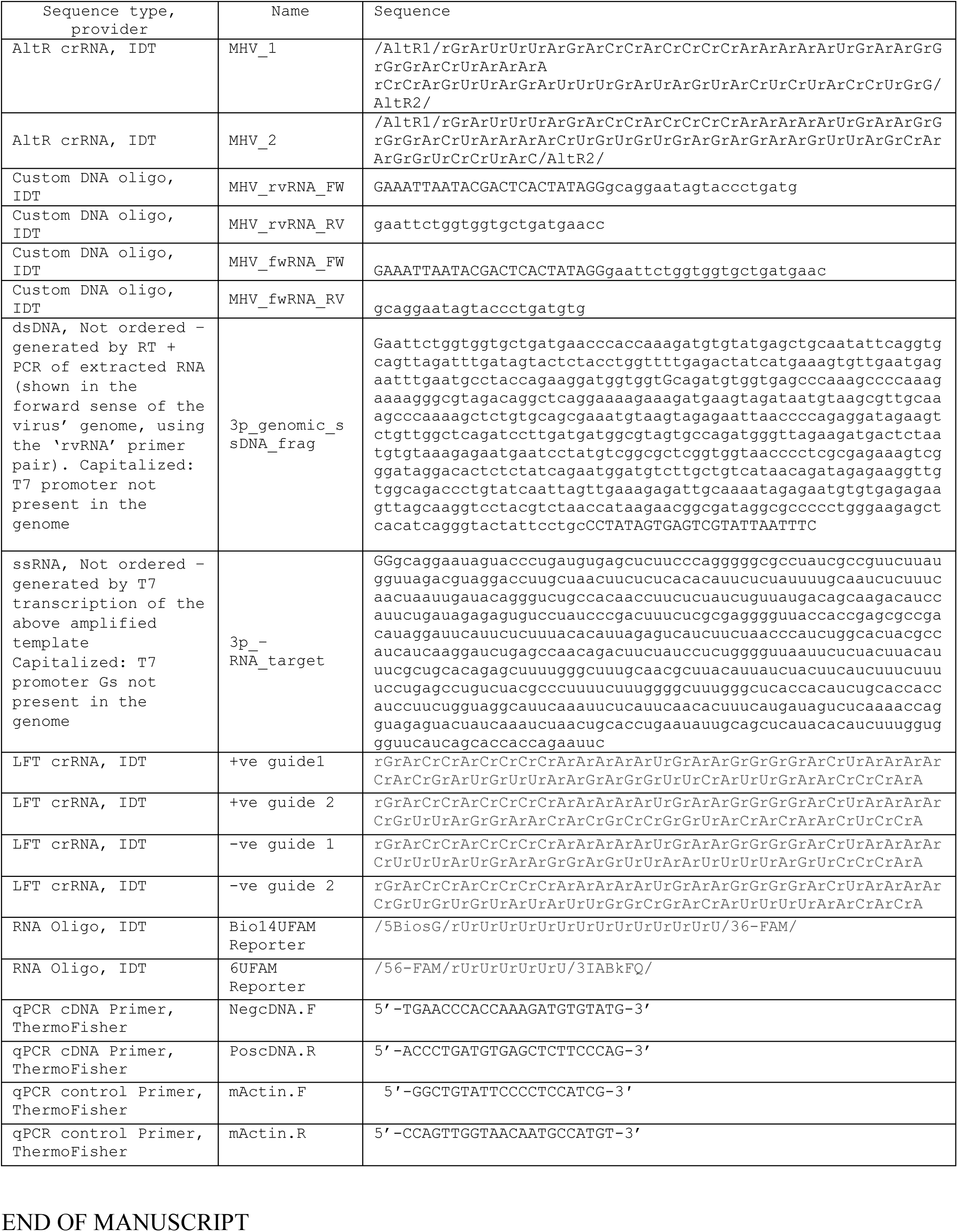
Sequences used in this study.

